# Human colon stem cells are the principal epithelial responders to bacterial antigens

**DOI:** 10.1101/2025.02.07.637053

**Authors:** Maaike H. de Vries, Claartje A. Meddens, Hemme J. Hijma, Anne-Claire Berrens, Suze A. Jansen, Berend A.P. Kooiman, Scott Snapper, Hans Clevers, Michal Mokry, Ewart W. Kuijk, Edward E.S. Nieuwenhuis

**Author notes:** Corresponding author: Lead contacts. Equal contribution (Ewart W. Kuijk, Edward E.S. Nieuwenhuis).

## Abstract

Intestinal epithelial cells (IECs) are capable of mounting an adequate antimicrobial inflammatory response to pathogens while tolerating commensals. The underlying regulatory mechanisms of immune sensitivity remain incompletely understood, particularly in the context of human IECs. To enhance our understanding of the immune response of IECs to bacterial epithelial barrier breach, we investigated whether epithelial responsiveness is contingent on cell identity and cell polarization. We exposed human intestinal organoids to bacterial antigens to study their immune responses. Notable discrepancies were observed in the specific reactions exhibited by intestinal stem cells (ISCs) and enterocytes. It was determined that basolateral exposure of IECs to bacterial antigens resulted in a robust response, whereas apical exposure elicited a significantly more modest response. We identified ISCs as the responders, while the reaction of enterocytes was found to be attenuated. The regulation of bacterial responsiveness in enterocytes occurs at multiple levels, including the modulation of NFκB activation and post-transcriptional control of mRNA stability. Our findings demonstrate that differentiated non-responsive enterocytes can be sensitized to bacterial antigens through the activation of the WNT pathway. These findings extend the crucial role of WNT signaling for intestinal epithelial homeostasis and regulation of stem cell maintenance, proliferation, differentiation, and tissue architecture in the gut. Additionally, they reveal a new function of WNT signaling in regulating microbial responses within the intestinal environment.

## Introduction

The large intestine is populated by a vast number of commensal microorganisms, resulting in a constant microbial exposure of the intestinal epithelial cells (IECs) [1, 2]. Under physiological conditions, IECs do not elicit an inflammatory response [3], however, inflammatory responses are necessary upon encountering pathogenic bacteria or during epithelial barrier disruption to prevent microbial invasion and dissemination into the bloodstream. Failure of IECs to maintain a balance between responsiveness and non-responsiveness during continuous microbial challenge [4, 5] can lead to chronic inflammation as observed in Inflammatory Bowel Disease (IBD).

The responsiveness of cells to microbes is dependent on the presence of Pattern Recognition Receptors (PRRs) and the expression and modification of molecules involved in intracellular signal transduction as well as various transcriptional and post-transcriptional processes [6, 7]. PRRs have the capacity to activate a multitude of immune response pathways, including the NFκB pathway [8]. In its inactive state, NFκB is bound to an inhibitor, IκB, which keeps it sequestered in the cytoplasm. Upon activation, IκB is phosphorylated and subsequently degraded, thereby releasing NFκB and enabling its entry into the nucleus. Within the nucleus, NFκB binds to specific DNA sequences in the promoter regions of target genes, thereby promoting transcription of NFκB target genes, including *CXCL8*, *TNFalpha*, and *NFKBIA* [8, 9]. The innate immune response requires minutes to hours to be fully activated upon stimulation [10].

PRRs are differentially expressed in the different epithelial cell types and along the apical basolateral axis, which may cause cell type specific responses [11–13]. All epithelial cell types originate from intestinal stem cells (ISCs) that reside at the bottom of the crypts. Differentiated progeny that emerge from these stem cells migrate towards the top of the villi of the small intestine or the inter-crypt regions of the colon, where the cells are shed into the lumen [14]. The self-renewal, proliferation and differentiation of ISCs are largely regulated by the WNT signaling pathway, which exhibits differential activity along the crypt-differentiation axis, with the highest activity observed in the crypt [15].

The precise relationship between differentiation state, responsiveness, and bacterial signaling remains unclear, as does the impact of bacterial signaling on cell identity [16–18]. While innate immunity pathways have been extensively studied in myeloid-derived cell types [19, 20], there is increasing interest for the role of for these pathways in IECs with most studies conducted in mouse models and immortalized cell lines. While these model systems provide valuable insights, each model has inherent limitations in studying immune responsiveness [21–23], such as intrinsic discrepancies between human and murine immune responses [21–25] and genetic aberrations that influence the responsiveness of human immortalized cell lines. Moreover, cell lines represent only a single cell type with incomplete cell differentiation and lack variation in genetic background [21–23, 26]. Human intestinal organoids can overcome these limitations, as they are derived from human ISCs isolated from epithelial biopsies and can be differentiated into various IEC types [27, 28]. Therefore, human intestinal organoids can be used to study the microbe interaction in an isolated setting [29–31].

The objective of this study is to investigate the immune responses in ISCs and differentiated IECs using human intestinal organoids exposed to *E. Coli* antigens. Our findings reveal disparities in immune responsiveness to bacterial antigens between ISCs and enterocytes, the most prevalent differentiated cell type of the intestinal epithelium. This study further explores the molecular mechanisms that underlies high responsiveness of ISCs and low responsiveness of enterocytes, dissecting the involved immune pathways at multiple levels, ranging from NFκB activation to post-transcriptional regulation of inflammatory gene transcripts. Our findings indicate that the intestinal immune response is, regulated processes upstream as well as downstream of NFκB activation with a governing role for the WNT pathway.

## Results

### 1. Epithelial cells mainly respond to bacterial antigens when exposed basolaterally and in a stem cell state

The intestinal epithelium, in conjunction with the mucosal layer, serves as the primary cellular defense mechanism against the gut microbiota [32, 33]. To investigate the responsiveness of IECs to microbial antigens, we utilized healthy, human colon-derived organoids and exposed them to a bacterial lysate. To assess the responsiveness of ISCs versus enterocytes, we conducted a comparative analysis of the effects of bacterial stimulation on undifferentiated stem cell-enriched organoids and differentiated enterocyte-enriched organoids followed by RNA-seq. Like the stem cells of the crypts, stem cell-enriched organoids have high WNT-pathway activity, while enterocyte organoids have low WNT-pathway activity. Enterocyte organoids were obtained from stem cell enriched organoids after five days of differentiation in the absence of WNT. Following a six-hour exposure to bacterial lysates, the undifferentiated organoids exhibited a strong response leading to the differential expression of over 1,000 genes (712 upregulated and 314 downregulated; **Figure 1A** and **Table S1**). This time point was used as a reference in all subsequent assays, unless otherwise indicated. The response is characterized by the upregulation of numerous inflammatory genes, including *TNFalpha*, *CXCL8*, *NFKB2*, and *NFKBIA*. Enterocyte organoids exhibited a weak response to bacterial lysate with less than 100 differentially expressed genes (61 upregulated and 17 downregulated) (**Figure 1B** and **Table S2**). The baseline expression levels of *CXCL8* demonstrated a comparable expression between unexposed stem cell-enriched organoids and enterocyte-enriched organoids (**Figure 1C**). This shows that the difference in *CXCL8* expression upon exposure cannot be explained by a higher baseline expression level in undifferentiated cells.

**Figure 1.**
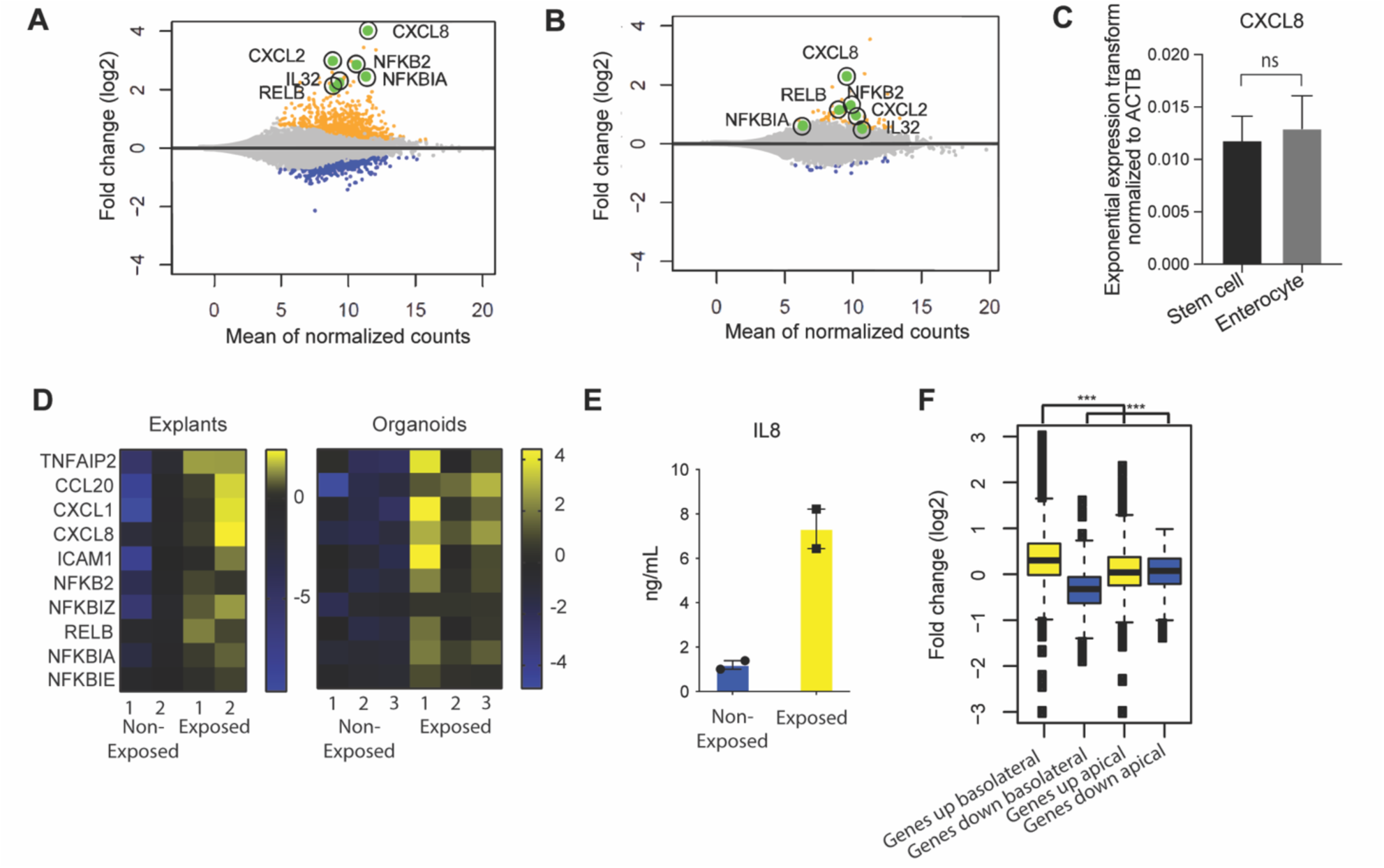
Epithelial cells mainly respond to bacterial antigens when exposed basolaterally and in a stem cell state. **A.** MA-plot-visualization of the log ratio and mean value of bulk RNAseq from healthy colon-derived stem cell-enriched organoids upon exposure to microbial antigens for 6h. (n=2). **B.** MA-plot visualization of the log ratio and mean value of bulk RNAseq from healthy colon-derived enterocyte-enriched organoids upon exposure to microbial antigens for 6h. (n=2). **C.** qPCR of CXCL8 of unexposed healthy colon-derived stem cell-enriched organoids and healthy colon-derived enterocyte-enriched organoids. Error bars indicate SEM (unpaired t-test) (n=3). **D.** Expression patterns of inflammatory genes of bulk RNAseq in healthy colon-derived stem cell-enriched organoids (exposed/non-exposed) compared to healthy colon explants (exposed/non-exposed). **E.** Luminex analysis of Interleukin-8 (IL8) release by healthy colon-derived stem cell-enriched organoids upon exposure to bacterial antigens 6h (n=2). Error bars indicate SEM. **F.** Expression changes of response genes were measured by bulk RNA-seq healthy colon-derived stem cell-enriched organoids grown as a monolayer and exposed from either the apical or basolateral side for 6h. Average genes up or down regulated. Statistical significance was determined using the Wilcoxon rank-sum test n=2. (results of the statistical test – as noted in the figure legend: ns = p > 0.05; * = p ≤ 0.05; ** = p ≤ 0.01; *** = p ≤ 0.001) qPCR = quantitative polymerase chain reaction.

Notably, we conducted a bulk RNA transcriptomics analysis of healthy colon explants (i.e., intact biopsies) and undifferentiated organoids exposed to bacterial lysates. The explants exhibited expression patterns analogous to those identified in the stem cell-enriched organoid model, indicating that the organoid experiments could be representative of responses from a completer tissue sample (**Figure 1D**).

Next, we analyzed the secretion of Interleukin-8 (IL8) protein when exposed to bacterial lysate. The colon stem cell-enriched organoids secreted IL8 upon exposure (**Figure 1E**). It can be concluded that the responses observed in the *in vitro* system reflect the intestinal epithelial inflammatory responses.

In physiological conditions, the intestinal epithelium is exposed exclusively to bacteria at the apical (luminal) surface only. To investigate the impact of cell polarity on microbial responsiveness, organoids were cultivated as monolayers that were exposed from the apical or the basolateral side. In contrast to the robust response observed upon basolateral exposure, apical responses were found to be attenuated under similar conditions (**Figure 1F, Figure S1)**. These findings suggest that in the *in vivo* situation, epithelial cells within the intestinal crypt only respond when the barrier that allows for microbial molecule passage is disrupted, resulting in basolateral exposure. This can occur when the barrier function of the intestinal epithelium is affected due to, for example, invasive microorganisms, genetic defects that are associated with increased epithelial permeability layer [34, 35] or disruption of the epithelium as observed in the inflamed intestinal mucosa of patients with IBD [36, 37]. Additionally, an uneven distribution of receptors along the apical-basolateral axis may contribute to the observed differences in apical and basolateral responsiveness [12, 13].

### 2. Single cell RNA sequencing shows that intestinal stem cells give a higher inflammatory response compared to enterocytes

To gain further insight into the specific responses of healthy human epithelial cell types during a breach of the epithelial barrier in their various states of differentiation, we performed single cell RNA sequencing (scRNAseq) on undifferentiated and differentiated colon organoid cultures exposed to bacterial lysate at the basolateral side. In this analysis, the stem cell and enterocyte conditions were pooled, resulting in a total of 911 cells. The data revealed the presence of seven distinct cell clusters (**Figure 2A**), which collectively represent the various stages of the three major cell types of IECs that are predominant under these culture conditions. These included stem cells, transit amplifying (TA) cells and enterocytes, which are found in their representative culture condition (**Figure S2**). The stem cells and the TA cells are composed of two clusters each. The distinctive feature that determines this separation is the cycling activity of cells, which is marked by the expression of *MKI67* (**Figure 2A, B, C**). Enterocyte-enriched organoids are separated into three clusters. One cluster consisted of enterocytes in the early phase of differentiation, based on the relatively low expression of differentiation markers. The other two clusters contained more differentiated enterocytes (**Figure 2A, B, C**).

**Figure 2.**
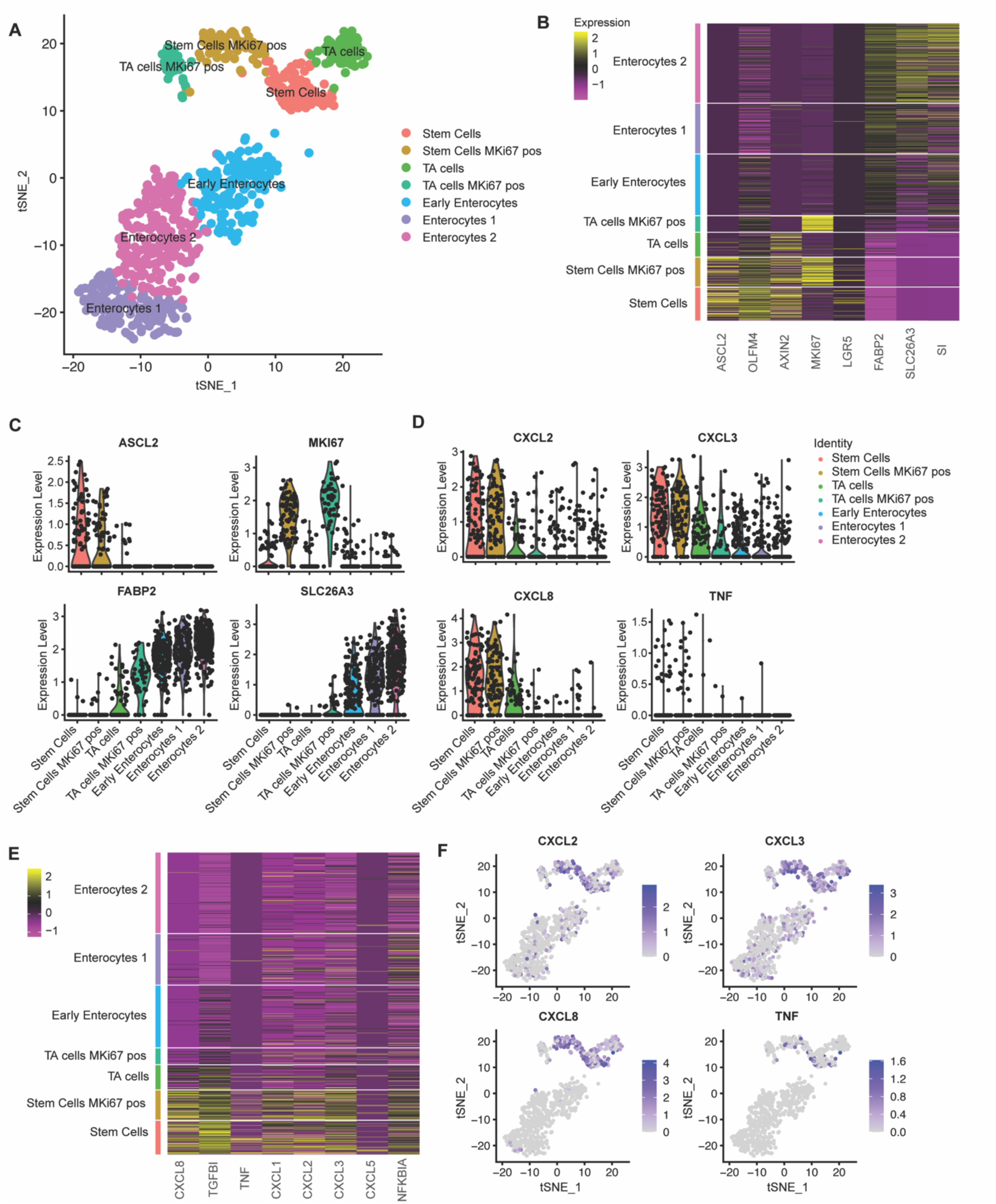
Single cell RNA sequencing shows that intestinal stem cells give a higher inflammatory response compared to enterocytes. **A.** tSNE-plot of cell clusters representing different subtypes of IECs. Data were pooled from five days differentiated enterocytes and stem cells colon-derived organoids upon exposure to 6h microbial antigens. **B.** Heatmap depicting the expression of stemness and differentiation markers in each identified cell type. **C.** Expression of markers that are associated with different states of epithelial differentiation, MKI67, SLC26A3, FABP2 and ASCL2 as determined per cell type. **D.** Expression of pro-inflammatory genes per cell type. **E.** Heatmap depicting the expression of inflammatory markers for each identified cell type **F.** Distribution of expression of CXCL2, 3, 8 and TNFalpha by tSNE-plots. TA = transit amplifying cells; pos = positive.

Subsequently, the expression levels of multiple bacterial response genes were projected onto the identified clusters. Cytokine expression was observed to be higher in the stem cell cluster in comparison to both the TA cell and the enterocyte clusters. A number of cytokines that are induced upon exposure to bacterial antigens were exclusively expressed within the stem cell clusters (**Figure 2D, E**). The upregulation of the inflammatory pathway was observed in all stem cells and was independent of *MKI67* expression (**Figure 2F**), thus excluding cycling activity as a crucial determinant for microbial responsiveness. These analyses confirm that, in comparison to enterocytes, stem cells are the inflammatory responders to microbial molecules among IECs in the large intestine.

### 3. The immune response of intestinal stem cell is NFκB-mediated

To gain a deeper comprehension of the consequences of bacterial stimulation, we conducted a gene set enrichment analysis on bulk RNA transcriptomics, which underwent alterations upon microbial exposure in ISCs. The enriched pathways exhibiting the most pronounced upregulation in undifferentiated organoids were predominantly associated with inflammatory responses (**Figure 3A**). It is noteworthy that a considerable number of immune response genes are regulated by the NFκB signaling pathway. Small molecule inhibition of NFκB during bacterial antigen exposure indeed resulted in reduced expression of the response gene *CXCL8* (**Figure 3B**).

**Figure 3:**
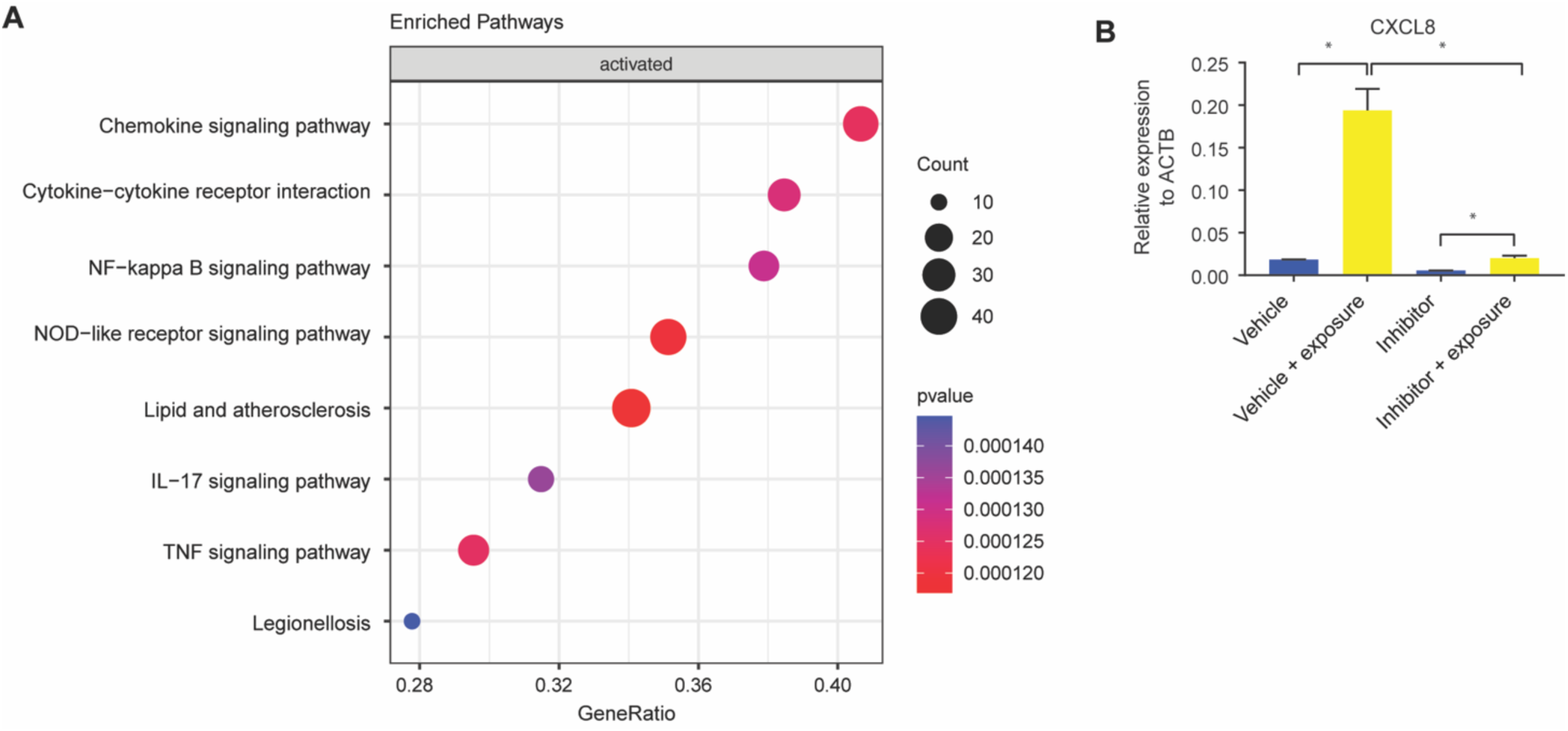
The immune response of intestinal stem cell is NFκB-mediated. **A.** Bulk RNAseq gene set enrichment analysis of upregulated genes of stem cell-enriched organoids. Response genes are enriched for inflammatory pathways. n=2. **B.** qPCR of CXCL8 of duodenal-derived stem cell-enriched organoid CXCL8 upon microbial exposure can be inhibited with a specific NFκB blocker. Error bars indicate SEM. (unpaired t-test) n=4.

### 4. Discrepancies in the expression of TLRs and the nuclear translocation of NFκB between stem cells and enterocytes

To elucidate the origin of the discrepancies in responses between stem cells and enterocytes, we conducted a comprehensive examination of the upstream and downstream activity of the NFκB pathway. The activation of NFκB is the result of a complex cascade of activating and inhibitory signals. Pathogen recognition molecules, such as Toll-Like receptors (TLRs), are activated by specific ligands [13]. Upon stimulation, these TLRs initiate a signaling cascade that leads to the phosphorylation and nuclear translocation of NFκB, ultimately activating innate immune response genes [38].

The differential expression of upstream NFκB components may be responsible for the observed differences in microbial responsiveness between stem cells and enterocytes. Our findings indicated that genes involved in the NFκB pathway exhibited higher expression levels in enterocyte-enriched organoids than in stem cell-enriched organoids. This suggests that the diminished responsiveness of enterocytes is not attributable to a global downregulation of this pathway (**Figure 4A, Figure S3**). Notable exceptions to this trend include *TLR5* and *TLR2*, both of which are expressed at higher levels in stem cell-enriched organoids than in enterocyte-enriched organoids. *TLR5* is activated by flagellin, while *TLR2* is activated by diverse range of microbial ligands [39]. Therefore, while the majority of NFκB pathway genes are upregulated during differentiation, the reduced expression of *TLR5* and *TLR2* in enterocytes may contribute to the diminished response to bacterial lysate. The differential expression of TLRs is consistent with previous findings in mice and pigs [11–13]. It is important to note, however, that there are inherent limitations to estimating pathway activity based on transcript abundance. It is not necessarily the case that there is a direct correlation between RNA transcript quantity and with protein translation [40]. Consequently, we also determined the nuclear localization of NFκB to examine the activation of the NFκB pathway in a downstream manner with respect to the TLRs. We conducted a comparative analysis of NFκB localization in stem cell- and enterocyte-enriched organoids after 2h of exposure to bacterial lysates. A higher In line with previous observations [41, 42], we observed NFκB translocation to the nucleus upon bacterial antigen exposure (**Figure 4B**, **4C**). It is noteworthy that nuclear NFκB translocation was highest in ISCs, but also evident in a substantial number of enterocytes, indicating that additional downstream mechanisms may further attenuate the responsiveness of enterocytes to bacterial molecules.

**Figure 4:**
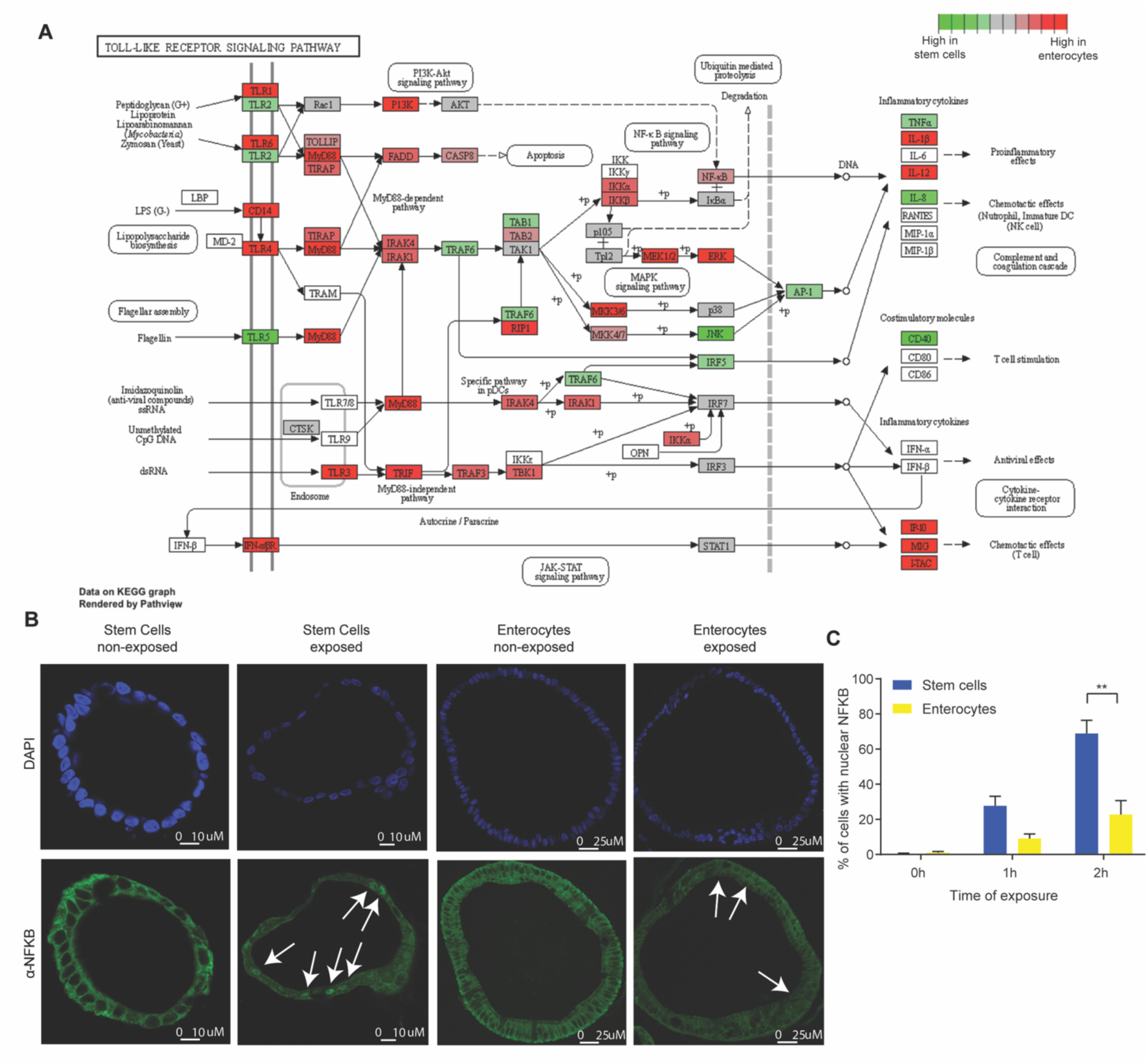
Discrepancies in the expression of TLRs and the nuclear translocation of NFκB between stem cells and enterocytes. **A**. Bulk RNAseq expression levels of genes involved in Toll-Like receptor signaling pathway in five days differentiated enterocytes-enriched organoids versus stem cell-enriched organoids. n=2 **B.** Immunohistochemistry of NFκB (p65) in stem cell-enriched organoids compared to five days differentiated enterocyte-enriched organoids, shows nuclear translocation of NFκB upon exposure (2h). n=3 **C.** Quantification of nuclear translocation of NFκB. The number of cells with nuclear NFκB is higher in stem cell-enriched organoids compared to enterocyte-enriched organoids (Mann-Whitney). Error bars indicate SEM. n=3 qPCR: quantitative polymerase chain reaction.

### 5. The chromatin in stem cells and enterocytes exhibits equal accessibility and activity of involved promoter sites

The subsequent analysis aimed to ascertain whether the chromatin accessibility and promoter activity of specific response genes contributes to the diminished microbial responsiveness observed in enterocytes. Chromatin immuno-precipitation (ChIP) was performed on colon organoids for histone modifications H3K4me3 and H3K27Ac, which are associated with active gene promoters [43]. It is notable that upon exposure to bacterial antigens, both stem cell- and enterocyte-enriched organoids exhibited the presence and induction of active chromatin marks within the promoters of the relevant response genes (**Table S3**), while the promoters of stem cell-associated genes were more active in stem cells and the promoters of differentiation-associated genes were more active in enterocytes (**Figure 5A**). For example, the promoter region of the differentiation marker *FABP2* is accessible in enterocytes and the promoter region of the stem cell marker *ASCL2* is accessible in undifferentiated cells. The promoter regions of the response genes *CXCL8* and *CXCL3* become accessible upon stimulation in enterocytes and stem cells (**Figure 5B**). These findings demonstrate that, even though the upregulation of the mRNAs of response genes is stronger in undifferentiated organoids, enterocyte-enriched organoids retain the ability to activate inflammatory gene promoters upon bacterial antigen exposure. This suggests that the reduced microbial responsiveness of enterocytes is not caused by decreased promoter accessibility.

**Figure 5:**
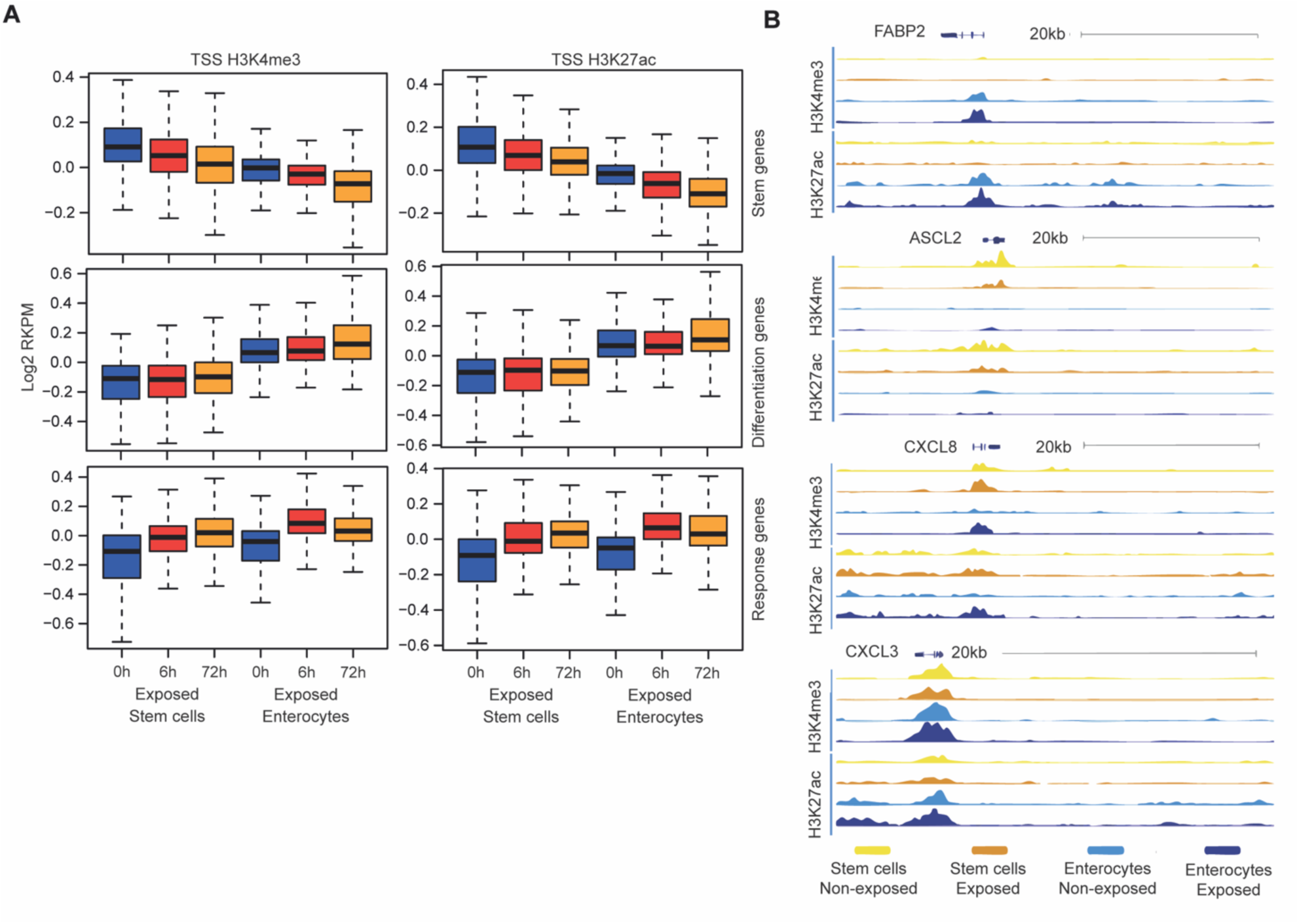
The chromatin in stem cells and enterocytes exhibits equal accessibility and activity of involved promoter sites. **A.** Chromatin immune-precipitation sequencing signal of active histone modifications (H3K4me3, H3K27ac, n=2) in the 2kb window around the transcriptional start site of different gene groups – at the baseline and upon exposure to bacterial antigens for 6h and 72h. (values represent median centered and log2 transformed RPKM) **B.** ChIP tracks at four genomic regions at baseline (-) and upon 6h of exposure to bacterial lysates (+) (n=2).

### 6. Evidence that post-transcriptional regulation may play a role in the half-life of genes involved in the epithelial inflammatory response

Prior research has indicated that post-transcriptional regulation may play a role in regulating immune response gene expression by facilitating the rapid degradation of inflammatory transcripts [44, 45]. For example, macrophage inflammatory responses are tightly regulated at the post-transcriptional level through AU-rich element (ARE)-mediated mRNA decay [46, 47]. This process involves the binding of RNA-binding proteins (RBPs) binding to AREs located in the 3’untranslated region (3’UTR) of mRNAs, which culminates in exonuclease-mediated degradation of the mRNA’s poly(A)-tail. This destabilization results in a reduction of the mRNA half-life and the prevention of translation. To ascertain whether AREs are present in IEC transcripts, we conducted an analysis to determine ARE enrichment in specific gene sets. The response genes (i.e. genes that are upregulated upon bacterial antigen exposure in stem cell-enriched organoids) exhibited a higher ARE content compared to the stem cell genes (stem genes are genes upregulated in stem cell-enriched organoids without exposure) and the differentiation genes (genes upregulated in enterocyte-enriched organoids without exposure) (**Figure 6A** and **Table S3**) [48]. This observation is consistent with previous studies that have reported the presence of AREs in approximately 22.4% of human mRNAs, particularly within the 3’UTR of inflammatory genes [48, 49]. This finding suggests that inflammatory genes in IECs are susceptible to post-transcriptional regulation [44, 50–52], which may play a role in the immune response of IECs.

**Figure 6.**
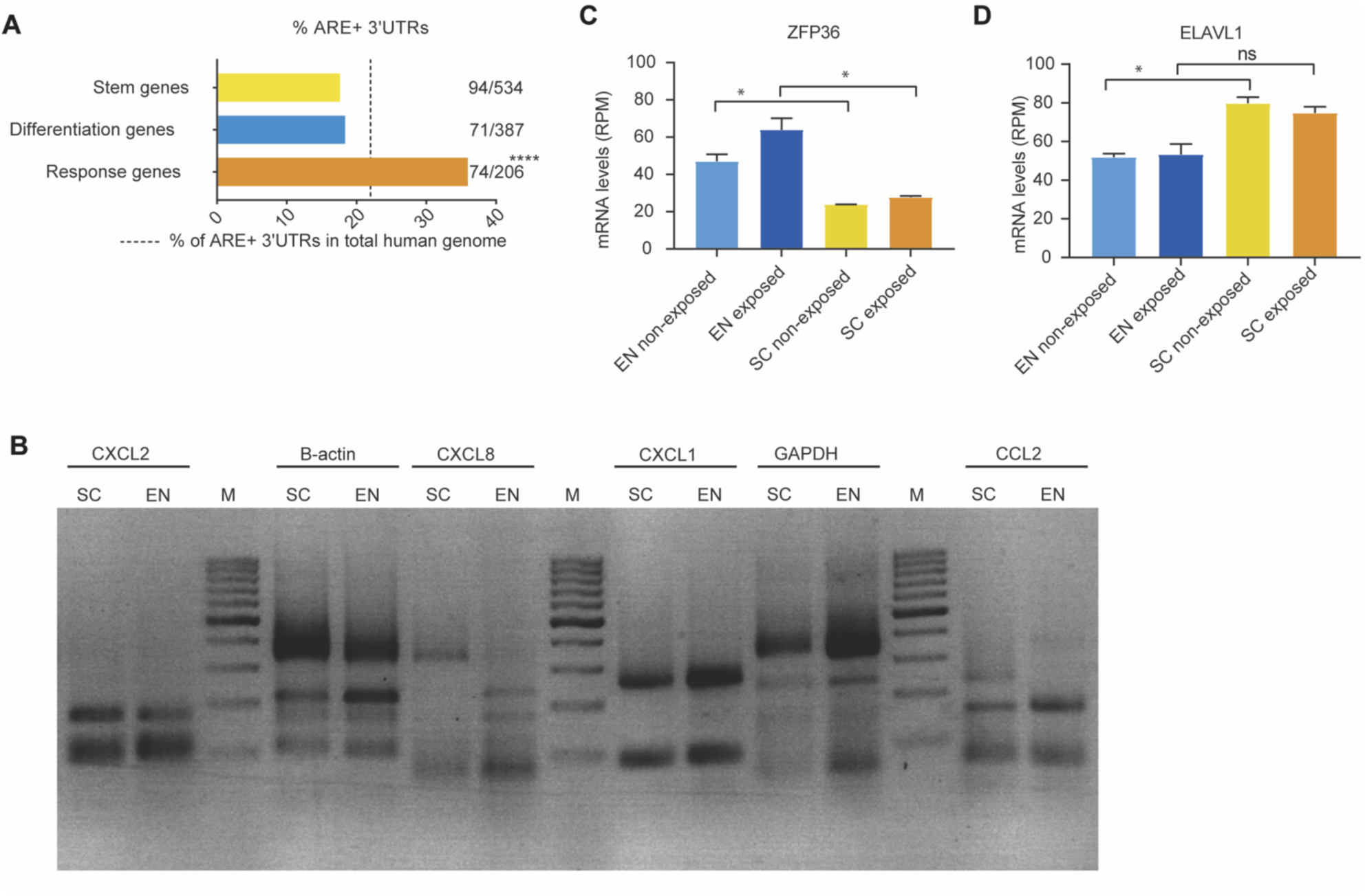
Evidence that post-transcriptional regulation may play a role in the half-life of genes involved in the epithelial inflammatory response. **A.** Presence of ARE-elements in 3’UTR of stemness, differentiation and response genes. (Fisher Exact test) n=2 **B**. poly(A) tail length assay 2% agarose gel. Bacterial exposure in stem cells-enriched organoids and enterocyte-enriched organoids for 6h. Marker is 100bp. M=Marker, EN=Enterocytes SC=Stem Cells. n=2 **C.** Bulk RNA expression of ZFP36 in non-exposed and 6h exposed colon-derived organoids. Error bars indicate SEM. EN=Enterocytes SC=Stem Cells. n=2 **D.** Bulk RNA expression of ELAVL1 in non-exposed and 6h exposed organoids. Error bars indicate SEM. (unpaired t-test) EN=Enterocytes SC=Stem Cells. n=2

The degradation of the poly(A)-tail represents a pivotal step in the pathway of mRNA decay, thus serving as a key determinant of mRNA half-life [53]. To explore whether the attenuated inflammatory response in enterocytes could be attributed to increased mRNA decay, we assessed the poly(A)-tail length of transcripts in our organoid models. To measure the poly(A)-tail length, we employed a method involving the addition of guanosine and inosine to the poly(A)-tail, followed by reverse transcription using a 3’UTR-specific primer and a universal reverse primer to generate complementary DNA. Subsequently, the amplified DNA was then analyzed by PCR, and product lengths were determined using agarose gel electrophoresis. The poly(A)-tail length of housekeeping genes (*ACTB* and *GAPDH*) and immune response genes (*CXCL2*, *CXCL8*, *CXCL1*, and *CCL2*) was assessed in stem cell-enriched and enterocyte-enriched organoids exposed to bacterial antigens. The poly(A)-tail length of housekeeping genes remains consistent in both stem cell state and enterocyte state. However, immune response genes exhibit either a similar length (*CXCL1* and *CXCL2*) or a shorter length in enterocyte-enriched samples (*CXCL8* and *CCL2*) compared to stem cell-enriched samples (**Figure 6B**). These findings suggest that a shorter poly(A)-tail length may contribute to the diminished inflammatory response of enterocytes, potentially due to faster mRNA degradation.

Previous studies have demonstrated that the RBP Zinc Finger Protein 36 (also known as TTP and encoded by the gene *ZFP36*) is capable of binding to AREs and serves as a pivotal mediator in poly(A)-mediated mRNA destabilization, particularly in the context of immune response genes [54]. We observe that *ZFP36* is higher expressed in enterocytes (**Figure 6C**) compared to stem cells. In contrast, *ELAVL1*, which encodes a protein (HuR) involved in mRNA stabilization [55], was found to be downregulated in enterocytes compared to stem cells (**Figure 6D**). Earlier studies have demonstrated that the expression of immune response genes and *ZFP36* are regulated by NFκB. As such, activated TTP inhibits nuclear translocation of NFκB and downregulates the immune response [56, 57]. Our results suggest that HuR and TTP may serve as potential factors that determine the mRNA stability of inflammatory response genes, thereby contributing to the observed differences in responsiveness between the various epithelial cell types.

### 7. WNT activation re-establishes inflammatory responses in enterocytes

The WNT/β-catenin pathway plays a pivotal role in the proliferation and maintenance of ISCs in the crypt and as undifferentiated organoids [14, 15] and decreased activity of this pathway induces differentiation (**Figure 7A**). We observed that undifferentiated organoids with high WNT activity show a stronger response to bacterial antigens than enterocyte organoids with low WNT activity. This may suggest crosstalk between the innate immunity and WNT pathways.

**Figure 7:**
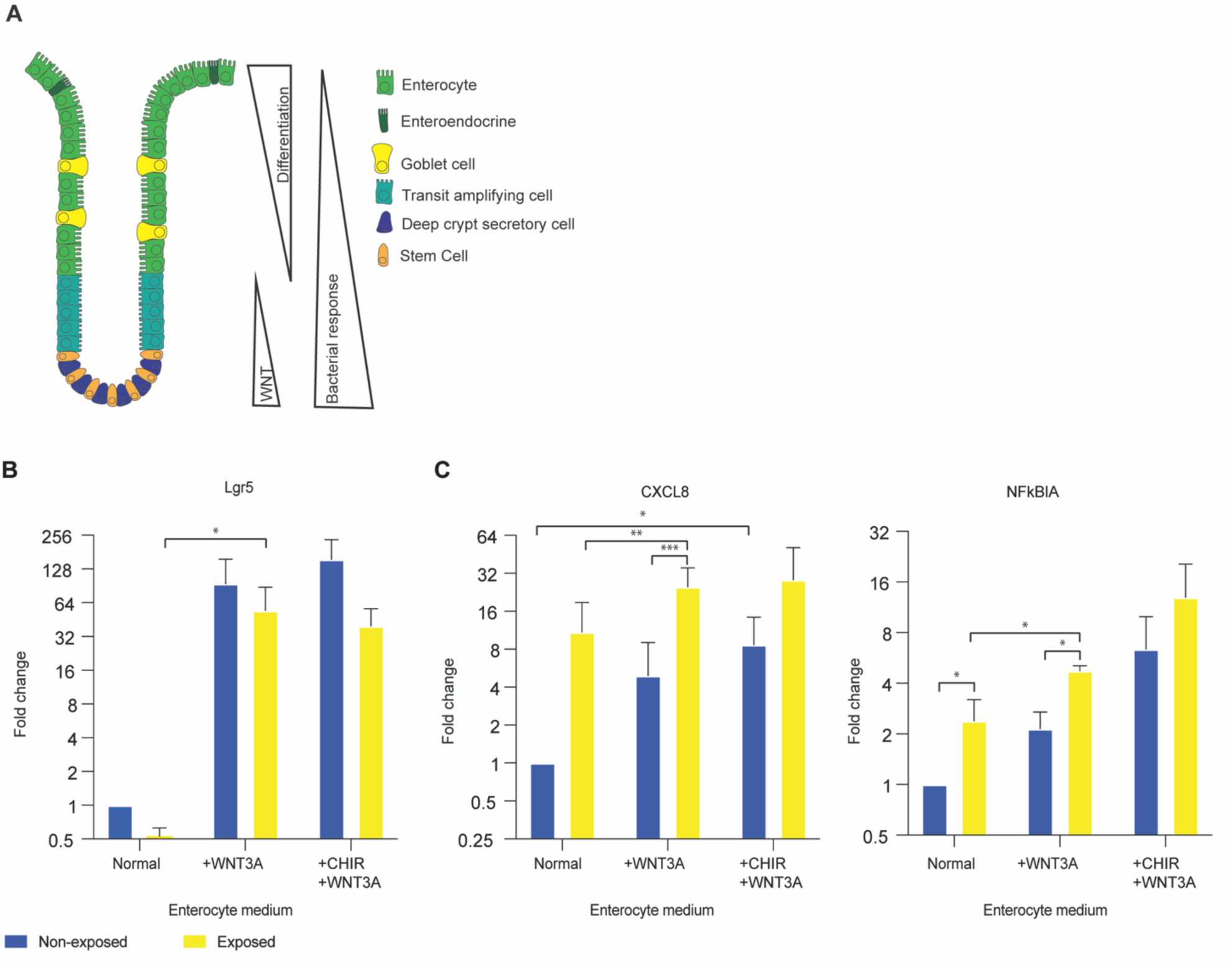
WNT activation re-establishes inflammatory responses in enterocytes. **A**. Transcription factor levels and microbial responsiveness along the crypt axis **B.** qPCR of LGR5 in enterocyte medium, enterocyte medium supplemented with 50% WNT3A conditioned medium, and enterocyte medium supplemented with 50% WNT3A conditioned medium and CHIR99021. Error bars indicate SEM (unpaired t-test) (n=3). **C**. qPCR of CXCL8 and NFKBIA in enterocyte medium, enterocyte medium supplemented with 50% WNT3A conditioned medium, and enterocyte medium supplemented with 50% WNT3A conditioned medium and CHIR99021. Error bars indicate SEM (unpaired t-test) (n=3). qPCR = quantitative polymerase chain reaction.

Therefore, we investigated if modulation of WNT signaling can determine the responsiveness of IECs. After 5 days of enterocyte differentiation in the absence of WNT, we stimulated the WNT pathway with WNT3A for a 24h period. While fully differentiated enterocytes do not have the plasticity to de-differentiate to stem cells [58], WNT-stimulation resulted in an increase in *LGR5* expression, confirming robust WNT-pathway activation (**Figure 7B**). Upon exposure to bacterial lysate in the presence of WNT3A, enterocyte-enriched organoids exhibited an increase in the expression of immune response genes *CXCL8* and *NFKBIA* (**Figure 7C).** Therefore, the observed increase in response genes suggests a direct involvement of the WNT pathway in the immune response to bacterial antigens.

The objective of the subsequent experiment was to ascertain whether the responsiveness of the stem cell-enriched organoids was diminished upon reduced WNT activity. To this aim, WNT3A was removed from the stem cell-enriched condition media to induce differentiation towards enterocytes. Following the withdrawal of WNT3A for a period of 24h, a slight decline in *LGR5* expression was observed (**Figure S4A**). When exposed to bacterial lysate in the absence of WNT3A, *CXCL8* and *NFKBIA* levels are comparable to those observed in stem cell-enriched organoids in the presence of WNT (**Figure S4B**). This suggests that a 24h period of WNT removal may not be sufficient to alter the stem cell state or its associated responsiveness to bacterial antigens.

To further assess whether increased WNT pathway activity modulates the responsiveness of stem cells, we introduced the WNT agonist CHIR99021 into the culture conditions alongside WNT3A for 24h [14]. CHIR99021 was administered to stem cell- and enterocyte-enriched organoids. This resulted in an increase in *LGR5* expression (**Figure 7B**, **Figure S4A**). Upon exposure to bacterial antigen, immune response genes *CXCL8* and *NFKBIA* were upregulated under these conditions (**Figure 7C**, **Figure S4B**). These results suggest that active WNT signaling enhances the NFκB-mediated immune response to bacterial exposure.

## Discussion

Our findings indicate that the ISCs are the responders to bacterial antigens in the large intestinal epithelium, in contrast to enterocytes. Moreover, our findings indicate that ISCs mainly respond to microbial stimuli when exposed basolaterally. These findings reinforce the mucosal paradigm of the intestinal epithelium, which is designed to avoid the elicitation of inflammatory responses to luminal antigens under homeostatic conditions. Conversely, inflammatory activation is only initiated when microbial antigens penetrate the IEC barrier and reach the basolateral surface of ISCs. The role of the ISCs as epithelial cell responders to microbial stimulation is consistent with previous reports that have identified these cells as having immune functions at the mucosal surfaces of the intestines. T-cell activation has been demonstrated to be contingent upon antigen presentation by ISCs via MHC-II expression [59]. Our study demonstrates that, in addition to antigen presentation, ISCs are involved in the mediation of local epithelial immune responses upon exposure to bacterial antigens. We illustrate that the variation in the response between ISCs and enterocytes to bacterial antigens is regulated at multiple levels upstream and downstream of NFκB activation. These findings highlight the intricate regulatory mechanisms that enable ISCs to function as immune mediators in the large intestine.

Lipopolysaccharides (LPS) are a widely utilized compound in the study of bacterial immune responses, as they represent the outer membrane components of gram-negative bacteria [60]. However, this approach offers a narrow view of the subject matter, as it does not encompass the full range of bacterial antigens present in more complex systems. In contrast, cecal slurry, prepared from the luminal contents of the colon, contains a more diverse range of bacterial antigens, including bacterial, viral antigens and food particles [61]. However, the presence of viral and dietary components in cecal slurry may complicate the analysis of a bacterial-specific immune response. Here we employed an *E. coli* lysate, which offers a controlled and diverse array of bacterial antigens while circumventing interference from non-bacterial components. This approach ensures a more targeted and reliable evaluation of bacterial immune responses.

Our observations indicated differential expression of TLRs and reduced nuclear NFκB translocation in enterocytes, suggesting differential regulation of the processes upstream of NFκB activation. Nevertheless, our results indicate that additional mechanisms may be involved in the diminished microbial responsiveness observed in enterocytes. It was thus established that enterocytes have shortened poly(A)-tails of specific inflammatory transcripts, which is likely to reduce the mRNA stability of these transcripts. It is postulated that mRNA stability in enterocytes is diminished by elevated activity of mRNA degradation pathways. This is corroborated by the observation of high expression of *ZFP36* and low expression of *ELAVL1*, which respectively exert a destabilizing and a stabilizing effect on mRNA. Regulation of the response to bacterial ligands may additionally extend to processes that determine translational and posttranslational activity.

Our findings indicate that in addition to its significant influence on the cell fate of IECs, the activity of the WNT pathway also plays a role in determining epithelial responsiveness. Prior research on other cell types and species has documented crosstalk between WNT/α-catenin signaling and NFκB [62–64]. Furthermore, α-catenin has been shown to bind to immune response genes on the DNA, thereby initiating their transcription [65]. The results of our study indicate that chromatin accessibility at promoter sites relevant to the inflammatory response is similar between stem cells and differentiated cells. These findings support the hypothesis that specific transcription factors may be directly involved in regulating microbial responsiveness.

The precise molecular mechanism through which WNT activity determines cellular responsiveness remains to be elucidated through further investigation. A potential explanation is through direct interaction between the key players of both pathways, α-catenin and NFκB, as has previously been described [62, 63]. Through such interaction, α-catenin could influence the nuclear translocation of NFκB upon stimulation with bacterial antigen, thereby promoting the NFκB-mediated immune response.

A number of genes are frequently mutated in IBD, and they play a role in the immune pathway. For example, there are mutations identified in the *TLR4*, *IL10R*, and *NFKBIZ* pathways [66–68]. The role of post-transcriptional regulation in the setting of inflammatory responses has been previously delineated in the context of immune cells [45]. The clinical relevance of these mechanisms is supported by the discovery of a genetic association between *ZFP36* family members (*ZFP36L1*, *ZFP36L2*) and IBD [69–71]. Furthermore, some patients with Very Early Onset-IBD have mutations in *TTC37*, which is involved in RNA decay [72, 73]. Other gene mutations involved in IBD affect IEC polarization [34, 35, 72]. Our data suggest that in cases where the epithelial barrier is compromised or the polarization is disrupted, resulting in exposure of the basolateral side is exposed to the microbiome, an inflammatory response may ensue.

The human organoid cultures lack the complete complexity of cell types present in the native epithelium [74]. To profile a heterogeneous population of cell types that constitute the majority of the human intestinal epithelium [75], we combined organoids grown under different culture conditions. In the future, models that include Paneth cells, goblet cells, tuft cells, or deep secretory cells [74] may provide a more comprehensive understanding of the differences in responsiveness among epithelial cells.

The advent of new techniques, such as whole exome sequencing and whole genome sequencing, has the potential to facilitate the discovery of additional genes that may be implicated in IBD. Moreover, the utilization of cutting-edge *in vitro* gut models will facilitate the elucidation of human gut-microbe interactions [30]. In light of the findings presented in this study, we anticipate that novel IBD-associated genes will be implicated in the processes upstream and downstream of NFκB and may be involved in pathways ranging from bacteria recognition to the post-transcriptional regulation of response genes, and WNT-signaling.

## Credit Authorship Contribution Statement

**MHdV**: Methodology, Formal analysis, Investigation, Writing - Original Draft, Writing - Review & Editing, Visualization. **CAM**: Methodology, Formal analysis, Investigation, Writing - Original Draft, Writing - Review & Editing, Visualization, Funding acquisition. **HJH**: Investigation. **ACB**: Investigation. **SAJ**: Investigation. **BAPK**: Investigation. **SS**: Resources, Funding acquisition. **HC**: Conceptualization, Funding acquisition. **MM**: Conceptualization, Methodology, Software, Writing - Review & Editing, Visualization, Supervision, Funding acquisition. **EWK**: Conceptualization, Methodology, Formal analysis, Writing - Review & Editing, Visualization, Supervision. **EESN**: Conceptualization, Methodology, Writing - Review & Editing, Supervision, Funding acquisition.

## Funding

MHdV, CAM, EWK, MM and EESN, were supported by the Leona M. and Harry B. Helmsley Charitable Trust., HC, and SBS were supported by NIH (RC2DK122532) Grant. MM was supported by the Career Development grant from MLDS (CDG-15). CAM was supported by the Alexandre Suerman stipend of the UMC Utrecht.

## Acknowledgements

We thank Dr. Sabine Middendorp for generating and providing the organoids and Prof. Dr. Sabine Fuchs for maintenance of the organoid biobank.

## Conflict of Interest

H.C. is the head of Pharma Research and Early Development at Roche, Basel, and holds several patents related to organoid technology. The full disclosure is given at https://www.uu.nl/staff/JCClevers.

## Data availability

Data will be made available upon publication.

## Abbreviations

ARE: AU-rich element
ChIP: chromatin immuno-precipitation
IBD: Inflammatory bowel disease
IL8: Interleukin-8
IECs: Intestinal epithelial cells
ISCs: Intestinal stem cells
LPS: Lipopolysaccharides
PRR: Pattern recognition receptors
qPCR: quantitative polymerase chain reaction
RBPs: RNA binding proteins
scRNAseq: Single cell RNA sequencing
TLRs: Toll like receptors
TA: Transit amplifying cells
UC: Ulcerative colitis
3’UTR: 3’untranslated region

## Materials and Methods

### Medical and Ethical Guidelines

Biopsies were obtained via ileo-colonoscopies and gastroscopies, which were conducted as part of the standard diagnostic procedure. Human Material Approval for this study was obtained from the Ethics Committee (Medisch Ethische Toetsings Commissie, METC) of the University Medical Center Utrecht (www.umcutrecht.nl/METC) or Boston Children’s Hospital. Experiments are performed with multiple donors.

### Organoid culturing

Biopsies of the colon and ileum were obtained via colonoscopy. The biopsies exhibited no macroscopic or pathological abnormalities. Crypt isolation and culture of human intestinal cells from biopsies were performed in accordance with the previously described methodology [28, 76]. The organoids were maintained long-term in a stem cell medium, which consisted of in Advanced medium/F12 (Gibco, 12634028) containing RSPO1, noggin, EGF (Peprotech, 315-09-1MG), A83-01 (Tocris Bioscience, 2939/10), nicotinamide (SIGMA-ALDRICH, N0636-500G), SB202190 (SIGMA-ALDRICH, S7067-25mg), and WNT3A. To induce differentiation, the cultures were maintained for 5-7 days in enterocyte medium, which is stem cell medium lacking nicotinamide, SB202190, and WNT3A. The conditioned media for RSPO1 (stably transfected RSPO1 HEK293T cells were kindly provided by Dr. C. J. Kuo, Department of Medicine, Stanford, CA), noggin, and WNT3A were utilized. The medium was refreshed every 2– 3 days and the organoids were passaged at a ratio of 1:4 approximately every 10 days (detailed media composition: **Table S4**).

Matrigel-embedded organoids (3D) were cultured in 70% Matrigel (BD Biosciences), diluted using growth factor-deficient medium (GF-), which consisted of Advanced DMEM supplemented with penicillin/streptomycin (GIBCO, 15140122), 1M HEPES (GIBCO, 15630080) and Glutamax 100x (GIBCO, 35050061). The primary intestinal organoids were cultured at 37°C, in 5% CO2. Intestinal epithelial monolayers (2D) were prepared according to the methodology previously described [77]. In brief, transwells (Corning Costar, Tewksbury, MA, USA) were coated with Matrigel (1:40 in PBS+ with Ca/Mg, Sigma-Aldrich, D8662-500ML) for 1 hour at RT. Subsequently, 2.5*10^5^ single cells were seeded on a transwell insert in the corresponding 24-well plate. A volume of 100µL and 600µL of medium was utilized in the apical and basolateral compartments, respectively. The monolayers were cultivated until they reached confluence, which was determent through microscopy and trans-epithelial electrical resistance (TEER) measurement. The primary intestinal monolayers were cultured at 37°C, in 5% CO2 environment.

### Exposure to Bacterial Antigens

A bacterial lysate was prepared from E. coli HST-08 Stellar competent cells (Takara Bio, 636766). The bacteria were subjected to a 20-min heat-inactivated process at 75°C, pooled in sterile PBS, and subsequently frozen for 1h at -80°C. Subsequently, the samples were subjected to sonicated using a Covaris ultrasonicator with the following settings: duty cycle of 20%; intensity of 10; cycles per burst of 500, and a total sonication time of 30 sec in a 13x65mm glass vial (Covaris), subjected to centrifugation at 10,000g for 30 min at 4°C, and subsequently filtered through a sterilizing filter. The quantity of bacterial lysate utilized for organoid exposure was determined through a titration process, ranging from 1µL to 20µL of lysate per 500µL medium. This was done to ascertain the concentration that elicits half of the maximum *CXCL8* response on mRNA in 3D cultures. Stem cell-enriched organoids were exposed at day 7-10, and enterocyte-enriched organoids were exposed after 5 days of differentiation. The same bacterial lysate concentrations were employed in 3D, and 2D apical and basolateral exposure experiments. In experiments involving multiple days of exposure, a new bacterial lysate was added at each medium refreshment (approx. every 2 days).

### RNA isolation and qPCR

RNA was isolated with TRIzol® LS (Ambion, cat. no. 10296-028), in accordance with the manufacturer’s protocol. cDNA was synthesized through reverse transcription (Invitrogen, Carlsbad, CA or iScript, Biorad, Hercules, CA, 1708891). Messenger RNA (mRNA) abundances were determined by real-time PCR using validated primer pairs and SYBR Green (Bio-Rad, Hercules, CA, 1708886). ACTB mRNA abundance was employed for normalization purposes. The following qPCR primers were utilized: *Lgr5* forward GAATCCCCTGCCCAGTCTC, *Lgr5* reverse ATTGAAGGCTTCGCAAATTCT, *α-actin* forward TGGCACCCAGCACAATGAA, *α-actin* reverse CTAAGTCATAGTCCGCCTAGAAGCA, *NFκBIA* forward GCAAAATCCTGACCTGGTGT, *NFκBIA* reverse GCTCGTCCTCTGTGAACTCC, *CXCL8* forward GGCACAAACTTTCAGAGACAG, *CXCL8* reverse ACACAGAGCTGCAGAAATCAG.

### Luminex

At the time of harvesting, the medium from organoids was collected (after 6h exposure) and was stored at - 80°C. The concentrations of CXCL8 were measured using the Luminex technology as previously described [78].

### RNA sequencing

RNA was isolated with TRIzol® LS (Ambion, cat. no. 10296-028), in accordance with the manufacturer’s protocol. Libraries were generated using NEXTflexTM Rapid RNA-seq Kit (Bio Scientific) and sequenced by the Nextseq500 platform (Illumina) to produce 75 bp single-end reads at the Utrecht DNA sequencing facility. Reads were aligned to the human reference genome GRCh37 using STAR. Differentially expressed genes in the transcriptome data were identified using the DESeq2 package with standard settings [79]. RPM values were calculated using edgeR’s RPKM function.

### Explants

Colon biopsies were washed and cultured in stem cell medium (described above) for 24h. Thereafter, the explants were stimulated with bacterial lysate for 6h. Next, RNA was then isolated and sequenced as described above.

### Pathway analysis, functional enrichment, GSEA, and KEGG pathway analysis

Functional enrichment was performed with the R-package clusterProfiler [80–82], while GSEA was carried out with the packages gage and gageData were used [83]. KEGG pathway analysis was performed using the KEGG pathway analysis package [84].

### Immunohistochemistry

The organoids were collected by meticulous disruption of the Matrigel and sequential elimination of the Matrigel through centrifugation (5 min, 2000rpm). The samples were subsequently fixed in 4% formaldehyde and embedded in 200µL of 2% agarose in dH2O, prior to embedding the samples in paraffin. Slides with a thickness of 5µm were deparaffinized and subjected to heat-mediated antigen retrieval was performed for a period of 20 min in a citrate antigen retrieval buffer with a pH of 6 (Sigma-Aldrich, C9999). The slides were blocked for 30 min in 5% BSA at RT and incubated ON at 4°C with primary antibodies (mouse α-NFκB p65 L8F6 1:50 (CST 6956S, Cell signaling Technology) in 5% BSA-PBS). Subsequently, the slides were incubated with secondary antibodies Alexa 488 donkey-anti-mouse (1:400 (A21202, Thermo Fisher Scientific)) for 1h a RT. Images were captured with a 63x objective on a Leica TCS SP8 X confocal microscope.

### NFκB-inhibition

Duodenum organoids were cultivated from single cells on stem cell medium. 8 Days after seeding, the organoids were treated with NFκB-inhibitors: 5µM IMD 0354 (Abcam, ab144823) or 10µM TPCA 1 (Abcam, ab145522). Following the addition of the inhibitors after12h of inhibitors, the organoids were exposed to bacterial lysate for 6h, after which RNA was isolated.

### CHIR99021

Intestinal organoids generated from single cells using stem cell medium. The organoids were cultured from single cells for 10-14 days, either on stem cell medium or by differentiating them into enterocytes for the 5 days on enterocyte medium. 3μM of CHIR99021 (Bio-Techne, 4423/10) [14, 85] were added 18h prior to exposure of bacterial lysate for 6h, after which RNA was isolated.

### Single-cell RNA sequencing

Colon and ileum-derived organoids were cultured for 10-14 days from single cells in 3 conditions: 1) on stem cell medium during the whole experiment, 2) stem cell medium was changed to enterocyte medium for the last 24 hours before harvesting, 3) stem cells medium was changed for the last 4 days before harvesting. For each condition half of the organoids were exposed for 6h with bacterial lysates before harvesting. Subsequently, cells were trypsinized and FACS-sorted using PI (Thermo Fisher Scientific - P3566) to eliminate dead cells into 384-well pre-indexed plates per condition. The plates were processed by Single Cell Discoveries as described previously using SORT-seq technology [86]. The data were analyzed using the Seurat V2.3.4. software package after excluding mitochondrial and ribosomal gene and a set of unreliably mapped genes (UGDH-AS1, PGM2P2, LOC100131257, MALAT1, KCNQ1OT1, PGM5P2, MAB21L3, EEF1A1) with these parameters: CreateSeuratObject(min.cells = 3, min.genes = 1500), NormalizeData(normalization.method = "LogNormalize", scale.factor = 10000), FindVariableGenes(mean.function = ExpMean, dispersion.function = LogVMR, x.low.cutoff = 0.0125, x.high.cutoff = 3, y.cutoff = 0.5), FindClusters(reduction.type = "pca", dims.use = 1:12, resolution = 0.6).

### ChIP sequencing

Chromatin immunoprecipitation was conducted using the MAGnify ChIP kit (Invitrogen, Carlsbad, CA) in accordance with the manufacturer’s instructions. 1µL α-acetylated histone 3 lysine 27 (H3K27ac) (ab4729; Abcam) or 1µL α-trimethylated lysine 4 (H3K4me3) (#39159; Active Motif) was utilized per immunoprecipitation. The captured DNA was purified using the ChIP DNA Clean & Concentrator kit (Zymo Research). The libraries were prepared using the NEXTflex™ Rapid DNA Sequencing Kit (Bioo Scientific). The samples were PCR amplified, checked for the proper size range and for the absence of adaptor dimers on a 2% agarose gel, and barcoded libraries were sequenced 75 bp single-end on Illumina NextSeq500 sequencer. The sequencing reads were mapped against the reference genome (hg19 assembly, NCBI37) using the BWA package (mem –t 7 –c 100 –M –R)42. Multiple reads mapping to the same location and strand were collapsed to single read and used for peak-calling. Peaks/regions were called using Cisgenome 2.043 (–e 150 -maxgap 200 –minlen 200).

### ARE-enrichment

ARE-enrichment was calculated using the ARED-plus database [48]. The percentage of response genes (genes upregulated upon 6h exposure to bacterial lysate), stem genes (genes upregulated in stem cell medium without exposure, **Table S3**) or differentiation genes (genes upregulated in enterocyte medium without exposure, **Table S3**) that have at least one AU-rich element encoded in the 3’UTR was calculated per gene set.

### Poly(A)-assay

Colon organoids were cultured from single cells for a period of 10-14 days, either in stem cell medium or organoids were differentiated for the 5 days on enterocyte medium. All organoids were subjected to bacterial lysate for 6h before RNA isolation, as previously described. GI-tailing and reverse transcription were done using the Poly(A)-tail length assay kit (ThermoFisher Scientific, # 764551KT) and according to the manufacturer’s protocol. Next, amplification of individual genes was done with a gene specific forward primer (*CXCL8* CTTGTCATTGCCAGCTGTGT, *GAPDH* CAACGAATTTGGCTACAGCA, *α-actin* ATCCTAAAAGCCACCCCACT, *CXCL1 GGCATACTGCCTTGTTTAATGG*, *CXCL2 CACAGTGTGTGGTCAACATTTCT*, *CCL2 GATACAGAGACTTGGGGAAATTG*) and a GI-tail reverse primer (Poly(A) tail length assay kit) for 35 cycles in Platinum™ PCR SuperMix High Fidelity (Invitrogen, # 12532016). Samples were analysed on a 2% agarose gel.

