## Supplementary material for "Human colon stem cells are the principal epithelial responders to bacterial antigens": Figure S

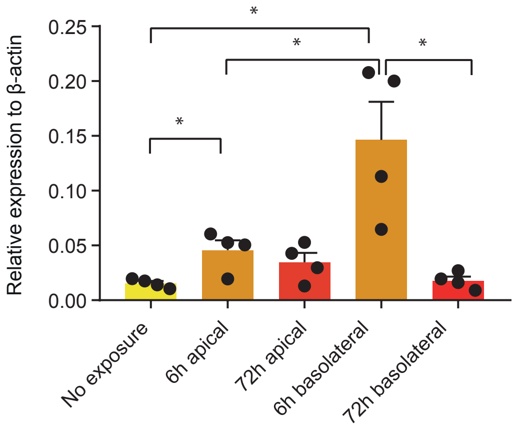


### Supplementary figure 1.

qPCR of CXCL8 on colon-derived organoids grown as monolayers upon apical and basolateral exposure to bacterial antigens (unpaired t-test). Error bars indicate SEM (n=4). qPCR = quantitative polymerase chain reaction

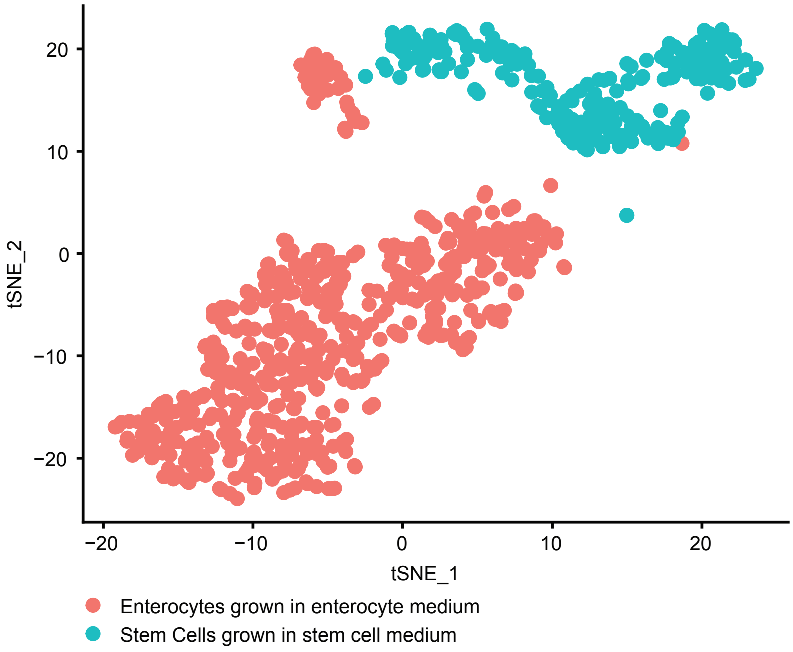


### Supplementary figure 2

Stem cell populations grown stem cell medium and enterocyte population grown in enterocyte medium.

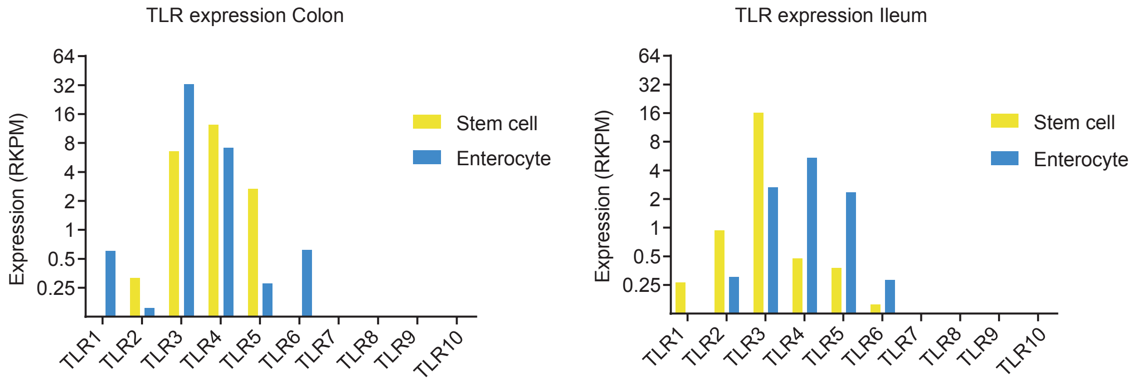


### Supplementary figure 3.

Bulk mRNA expression of Toll-like receptor (TLR) in enterocyte-enriched and stem cell-enriched colon-derived organoids.

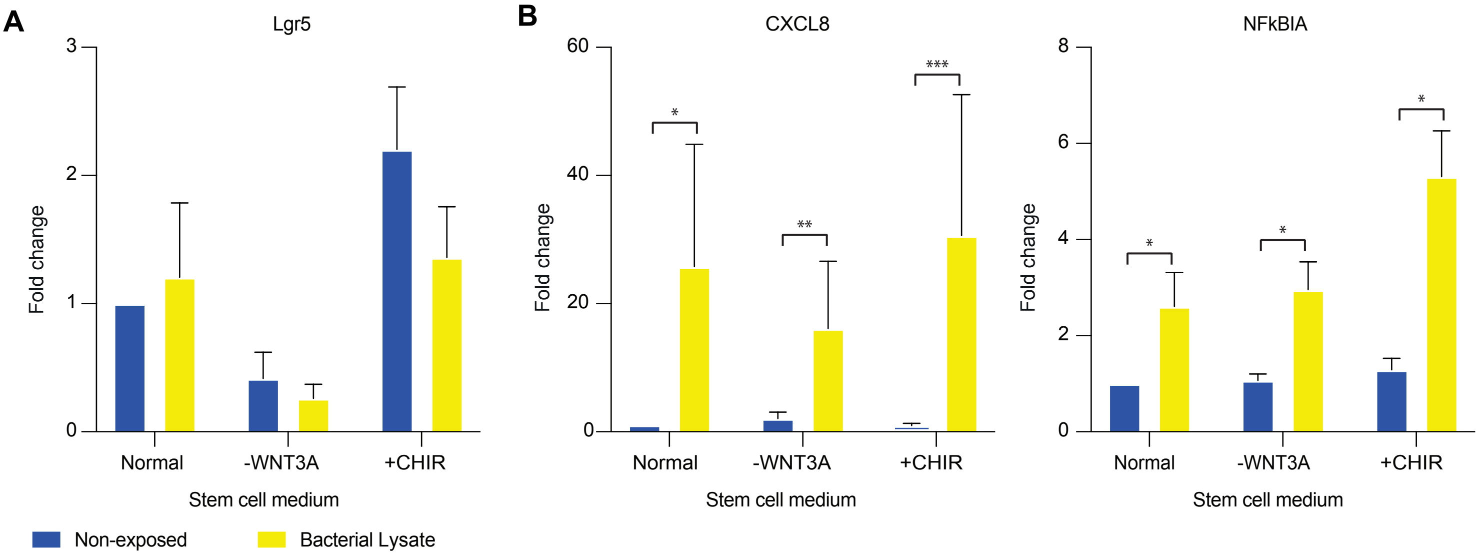


### Supplementary figure 4. WNT activation or deactivation and inflammatory responses in intestinal stem cells.

A. qPCR of Lgr5 in normal stem cell medium, stem cell medium without WNT3A conditioned medium, and stem cell medium with CHIR99021. Error bars indicate SEM (unpaired t-test) (n=3).

B. qPCR of CXCL8 and NFKBIA in normal stem cell medium, stem cell medium without WNT3A conditioned medium, and stem cell medium with CHIR99021. Error bars indicate SEM (unpaired t-test) (n=3).

qPCR = quantitative polymerase chain reaction.
